# ScreenCube: A 3D Printed System For Rapid and Cost-Effective Chemical Screening in Adult Zebrafish

**DOI:** 10.1101/163469

**Authors:** Adrian T. Monstad-Rios, Claire J. Watson, Ronald Y. Kwon

## Abstract

Phenotype-based small molecule screens in zebrafish embryos and larvae have been successful in accelerating pathway and therapeutic discovery for diverse biological processes. Yet, the application of chemical screens to adult physiologies has been relatively limited due to additional demands on cost, space, and labor associated with screens in adult animals. Here, we present a 3D printed system and methods for intermittent drug dosing that enable rapid and cost-effective chemical administration in adult zebrafish. Using pre-filled screening plates, the system enables dosing of 96 fish in ˜3min, with a tenfold-reduction in drug quantity compared to that used in previous chemical screens in adult zebrafish. We characterize water quality kinetics during immersion in the system, and use these kinetics to rationally design intermittent dosing regimens that result in 100% fish survival. As a demonstration of system fidelity, we show the potential to identify two known chemical inhibitors of adult tail fin regeneration, cyclopamine and dorsomorphin. By developing methods for rapid and cost-effective chemical administration in adult zebrafish, this study expands the potential for small molecule discovery in post-embryonic models of development, disease, and regeneration.

## INTRODUCTION

Zebrafish are a powerful model system for human disease modeling due to their amenability to experimental approaches to therapeutic discovery that are challenging in other vertebrate systems. A prime example of such an approach is phenotype-based small molecule screening. Most commonly, zebrafish embryos or larvae are loaded into the wells of a 96-well plate (typically filled with ˜200µL of solution/well ^1^), compounds from chemical libraries are administered to the water (upon which they are absorbed through the skin, gills, and/or digestive tract), and the effects on phenotype are measured. Such screens have been used to study diverse processes, tissues, and organs, mostly within the first few days of development (for comprehensive reviews, see^2, 3^).

Despite the fact that zebrafish from later stages of development (i.e., juvenile to adult) are being increasingly employed to study aspects of postembryonic development ^4^ ^5^ ^6^, disease ^7^, and regeneration ^8^ ^9^ ^10^, chemical screens in adults have been relatively limited due to their additional demands on cost, space, and labor. For instance, adult fish require a large volume of water for housing, increasing the quantity of drug necessary to achieve an active concentration in the water. In 2010, Oppedal and Goldsmith performed the first instance of phenotype-based small molecule discovery in adult zebrafish by performing a chemical screen for modulators of adult tail fin regeneration ^11^. To reduce the quantity of drug required to achieve an active concentration in the water, fish were housed in 100ml of water in 250mL specimen cups. However, even in this reduced volume, administering compound for three consecutive days at 5µM (with a 10mM stock solution provided in a chemical library) would require 250µL of stock solution, exceeding the quantity of drug supplied in many commercial compound libraries (100µL of 10mM stock solution). Since the fish were housed statically, the water and drugs were replenished daily to maintain water quality. These water exchanges can become time consuming when conducted on a large scale. While alternative methods for drug delivery in adult zebrafish such as oral gavage ^12^ and injection ^13^ reduce the amount of compound needed for administration, they require multiple steps to perform (including preparation of equipment, anesthetization, individual administrations, and recovery from anesthesia) and are therefore inefficient in terms of time and throughput. Thus, there is a need for higher throughput, more cost-effective drug administration methods to increase the efficiency and scalability of chemical screening in adult zebrafish.

Here we present ScreenCube, a 3D printed chemical screening system that enables rapid and cost-effective chemical administration in adult zebrafish. We are able to transfer 96 fish into pre-filled screening plates for dosing in ˜3min. This low-volume dosing in the screening plates allows for a tenfold-reduction in drug quantity compared to that used in previous chemical screens in adult zebrafish. We characterize water quality kinetics during intermittent low-volume immersion, and show that these kinetics may be used to rationally design intermittent dosing regimens that result in 100% fish survival. As a showcase of system fidelity, we demonstrate the potential to identify two chemical inhibitors of adult tail fin regeneration, cyclopamine and dorsomorphin.

## MATERIALS AND METHODS

### Zebrafish Care

All animal studies were approved by the Institutional Animal Care and Use Committee at the University of Washington. Wildtype zebrafish (Aquatic Research Organisms, Hampton, NH) ^10^ were housed in 28°C water on a 14:10h light:dark photoperiod using a commercial housing rack system (Aquaneering, San Diego, CA). Fish were fed commercial zebrafish diet (GEMMA Micro, Skretting, Stavanger, Norway) according to manufacturer’s recommendations. All fish used for the study were mixed sex adult animals of ˜30-35mm standard length (S.L.).

### 3D Printing

CAD files were generated in Pro/ENGINEER (PTC, Needham, MA) or FreeCAD (http://freecadweb.org/). For 3D printing, CAD designs were converted to STL files, uploaded to Makerbot 3D printing software, and printed using a Makerbot Replicator 2 desktop 3D printer in clear PLA plastic. All CAD and STL files are included as supplementary files (Supplementary Files 1-8).

### Fin Regeneration

Fin regeneration studies were performed as previously described ^10^. Briefly, zebrafish were anesthetized in 0.02% MS-222 (E10521; Sigma-Aldrich, St. Louis, MO) and subjected to 50% tail fin amputation using a straight razor blade. For *in vivo* imaging, fish were anesthetized in MS-222 and imaged under a brightfield stereomicroscope. Percent area regrowth was calculated as described previously ^10^.

### Chemical Dosing

Cyclopamine (item#: 11321, Cayman Chemical Company) and dorsomorphin (item#: 11967, Cayman Chemical Company) were dissolved in ethanol and DMSO (respectively) to form a 10mM stock solution. For chemical dosing, screening plates were filled with system water at a volume of 10mL/well, an appropriate volume of stock solution (10mM) was added to each well, stirred briefly, and the inserts (with fish inside) were quickly transferred into the plates. Plates were placed in a 28.5°C incubator during dosing. Following dosing, the inserts were removed from the screening plates, quickly rinsed in a 6L tank filled with system water to remove residual chemicals, and returned to housing. Unless otherwise noted, chemicals and water in the screening plate were replenished daily.

### Water Quality

Dissolved oxygen within the water of the screening plate was measured using a Pinpoint II Oxygen Monitor (American Marine, Ridgefield, CT) according to manufacturer’s recommendations. Total ammonia was measured using a colorimetric test kit (API, Chalfont, PA). Volumes were ratiometrically scaled down from manufacturer’s recommendations to minimize the volume of water required for sampling.

### Statistical Analysis

All statistical analyses were performed in Prism (Graphpad Software, La Jolla, CA). For analysis of water quality kinetics, instances of exponential decay were modeled using a standard one-phase decay equation as Y=Plateau+(Y0-Plateau)^*^exp(-K^*^time), and comparisons of models was performed using t-tests of fitted model parameters. All other cases of water quality kinetics were modeled using linear regression, and comparisons of linear regressions were performed using ANCOVA-based methods ^14^. For dose response studies, we performed one-way ANOVA followed by Fisher’s PLSD post-hoc test. Time-dependent effects of drug treatment were assessed via a two-way ANOVA with time and treatment as factors. p?<?0.05 was considered statistically significant. All data are presented as mean±SEM unless otherwise noted.

## RESULTS

### A Dual-Compartment System For Chemical Screening in Adult Zebrafish

We designed ScreenCube, a multi-component screening system (Fig 1A-1D) comprised of dual-compartment housing inserts with lids, custom 6-well screening plates, and insert holders. To facilitate transfer of inserts, we designed 6-well insert holders that hold inserts in a 2x3 array. The insert holders can be used to nest inserts in either 6L tanks (for housing on standard commercial recirculating systems) (Fig 1E), or a dedicated recirculating screening rack (Fig 1F). By housing fish on a recirculating system, we alleviated the need for daily water changes to maintain water quality, such as would be required if fish were housed statically in specimen cups. Each housing insert holds a single adult fish, and consists of a large-volume upper compartment that tapers into a small-volume, mesh-bottomed lower compartment. The inserts are nested in an outer tank such that the waterline is near the top of the insert, allowing for free movement of fish during normal housing (Fig 2A, 2B). During chemical dosing, the inserts are removed from the outer tank, causing the fish to quickly drain into the lower compartment. The inserts are quickly placed into the 6-well screening plates that have been prefilled with water (10mL/well) plus the appropriate compound (Fig 2A', 2B'), enabling drug absorption. In general, fish drain into the lower compartment with extremely high success; in the rare occassion that a fish adheres to the side of the insert, this can be resolved by lightly tapping on the insert wall. Once immersed in the screening plate, the fish quickly cessate movement.

**Figure 1:**
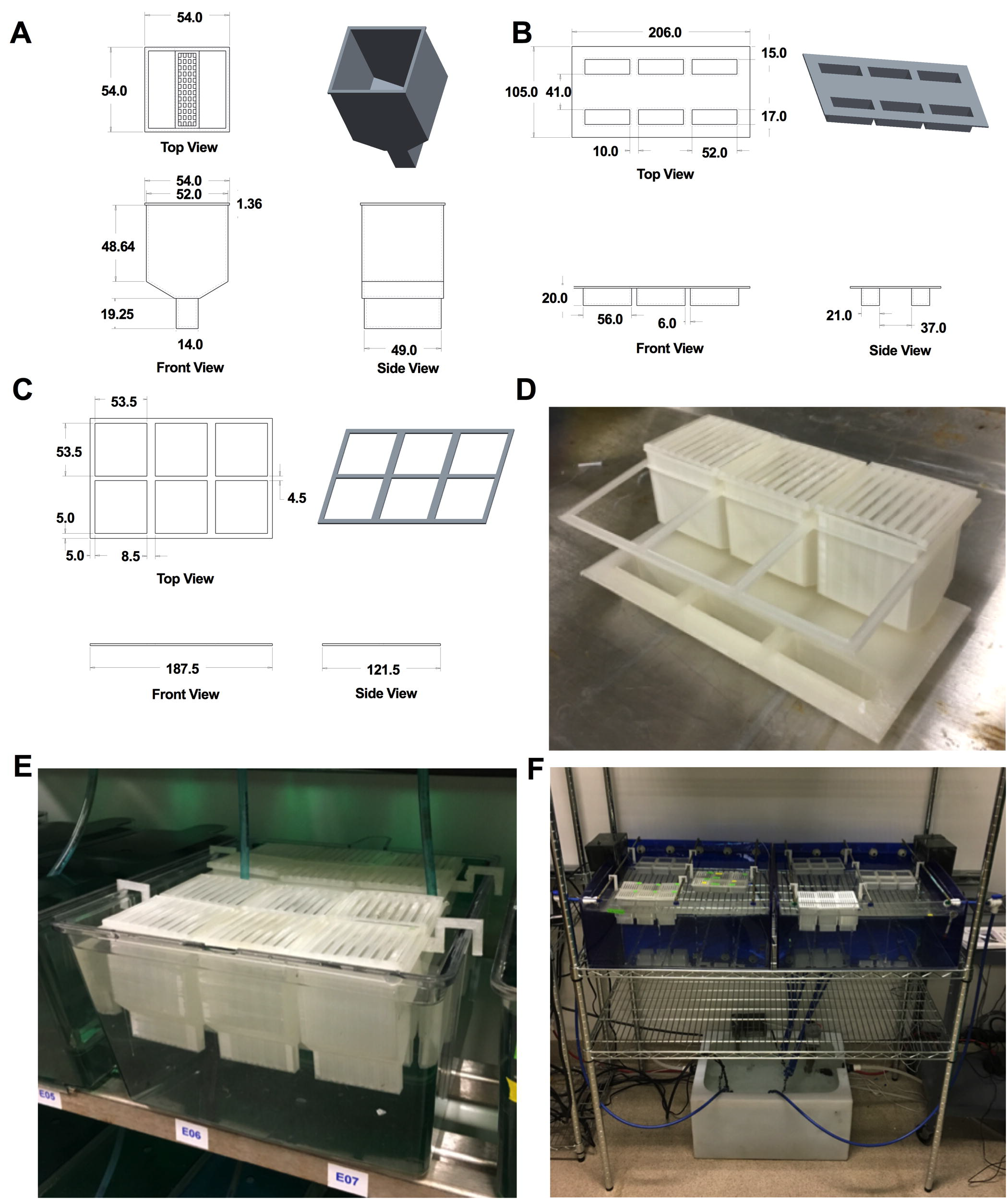
Design of a 3D printed system for chemical screening in adult zebrafish. (A-C): Mechanical drawings for the dual-compartment housing insert (A), 6-well screening plate (B), and insert holder (C). All dimensions are in mm. (D) Image depicting system components set up for drug administration (only three housing inserts are shown to enable view of the wells). (E) Housing inserts nested within a standard 6L tank on a commercial recirculating rack system. (F) Housing inserts nested within a custom screening rack. A benefit of a dedicated screening rack is that it reduces the potential for introducing residual compounds into the main system. Up to 12x insert arrays (each holding 6 inserts) can be housed in this system (only 6x are shown).

**Figure 2:**
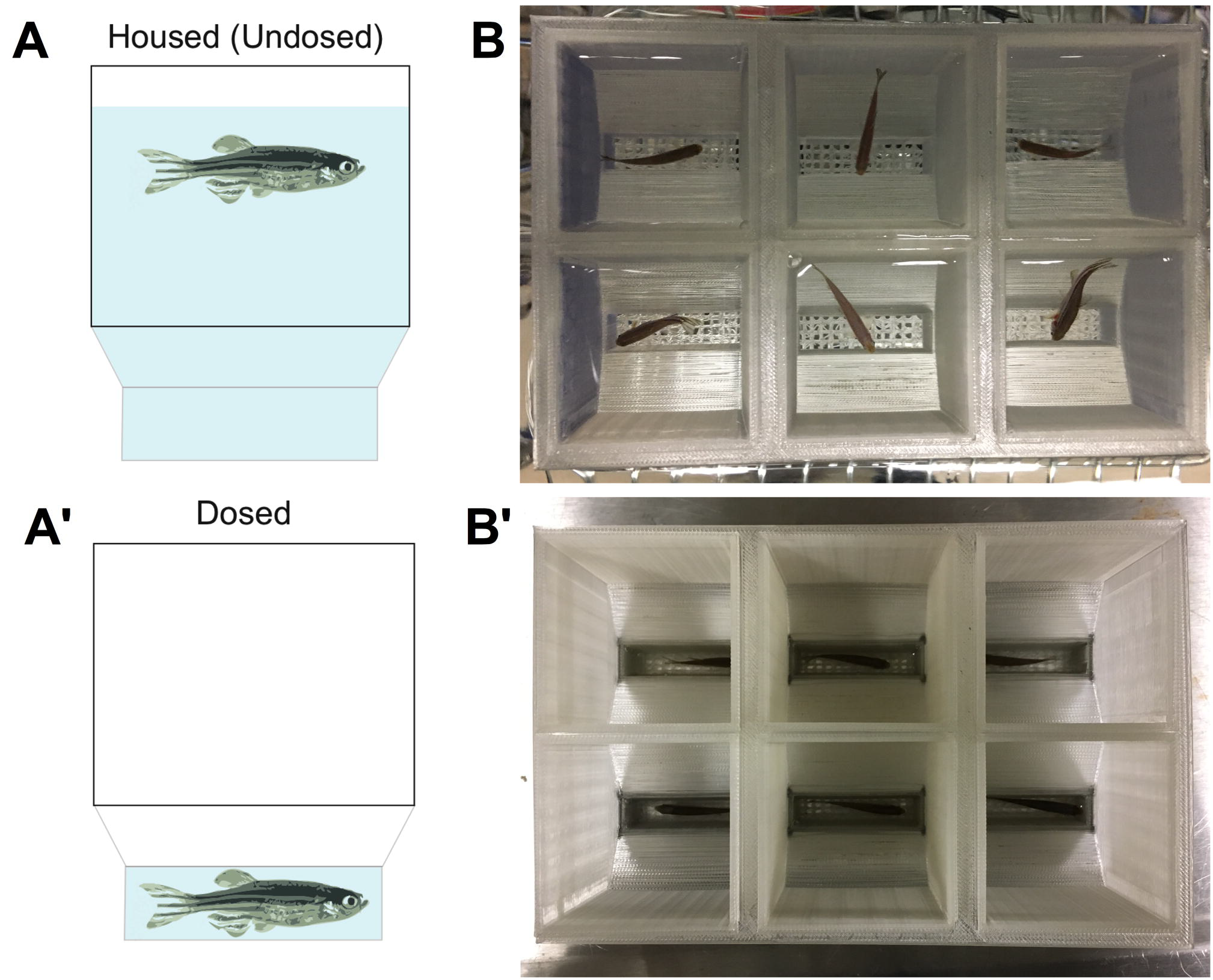
Mechanics of drug administration. (A, B) Each housing insert consists of a large-volume upper compartment that tapers into a small-volume, mesh-bottomed lower compartment. During normal housing, the inserts are nested in an outer tank such that the waterline is near the top of the insert (A', B') During chemical dosing, the inserts are removed from the outer tank, and quickly placed into pre-filled screening plates. The fish quickly drain into the lower compartment and cessate movement following immersion.

### Kinetics of Water Quality

Due to the small volume of water within the wells of the screening plate, the duration in which zebrafish may be safely dosed during intermittent low-volume immersion is dictated by changes in water quality. Sublethal uniodized ammonia poisoning in zebrafish can occur at levels as low as 0.02ppm ^16^. For the water temperature (28°C) and pH (7.2) employed in this study, the percentage of total ammonia present in unionized form is ~1% ^16^, suggesting that sub lethal poisoning can occur at total ammonia levels of 2ppm. The dissolved oxygen requirements of adult zebrafish have not been systematically determined ^17^. However, dissolved oxygen levels <3ppm are known to be stressful to most aquatic organisms ^18^.

We characterized dissolved oxygen and total ammonia kinetics while fish were immersed in the screening plate wells (Fig 3). In fish fed both 1h and 24h prior to immersion, dissolved oxygen decayed exponentially, with most changes occurring in the first twenty minutes (Fig 3A). When fit to a standard one-phase decay model, we observed a significantly higher decay constant and significantly lower plateau level in fish fed 1h prior to experimentation compared to those fed 24h prior (Table 1). The increased oxygen consumption in fish fed 1h prior is consistent with the notion that oxygen consumption in fish increases following food ingestion due to increased metabolic activity ^15^. In regard to total ammonia, we observed a linear increase in concentration with time (Fig 3B), with the slope significantly greater in fish fed 1h prior compared to those fed 24h before (Table 2). Thus, feeding of fish within an hour of dosing impacts water quality by both altering fish oxygen consumption as well as ammonia secretion. Our studies suggest that the time in which a fish may be continuously dosed within a single dosing bout is limited by the decay in dissolved oxygen, rather than accumulation of total ammonia. Specifically, for fish fed either 1h or 24h before dosing, the time required for dissolved oxygen to decay to 3ppm is less than the time required for total ammonia to accumulate to 2ppm.

**Table 1:** Exponential decay model parameters describing the kinetics of dissolved oxygen within a single dosing bout.

| Parameter | 24 Hours Prior | 1 Hour Prior | Comparison |
| --- | --- | --- | --- |
| Y0 (ppm) | 8.00±0.21 | 8.00±0.10 | p>0.99 |
| Plateau (ppm) | 2.56±0.08 | 2.25±0.04 | p=0.001 |
| K (1/min) | 0.07±0.01 | 0.11±0.01 | p=0.0005 |
| R square | 0.96 | 0.99 |  |

**Table 2:** Linear regression model parameters describing the kinetics of total ammonia within a single dosing bout.

| Parameter | 24 Hours Prior | 1 Hour Prior | Comparison |
| --- | --- | --- | --- |
| Slope (ppm/hour) | 0.55±0.02 | 1.71±0.06 | p<0.0001 |
| R square | 0.96 | 0.99 |  |

**Figure 3:**
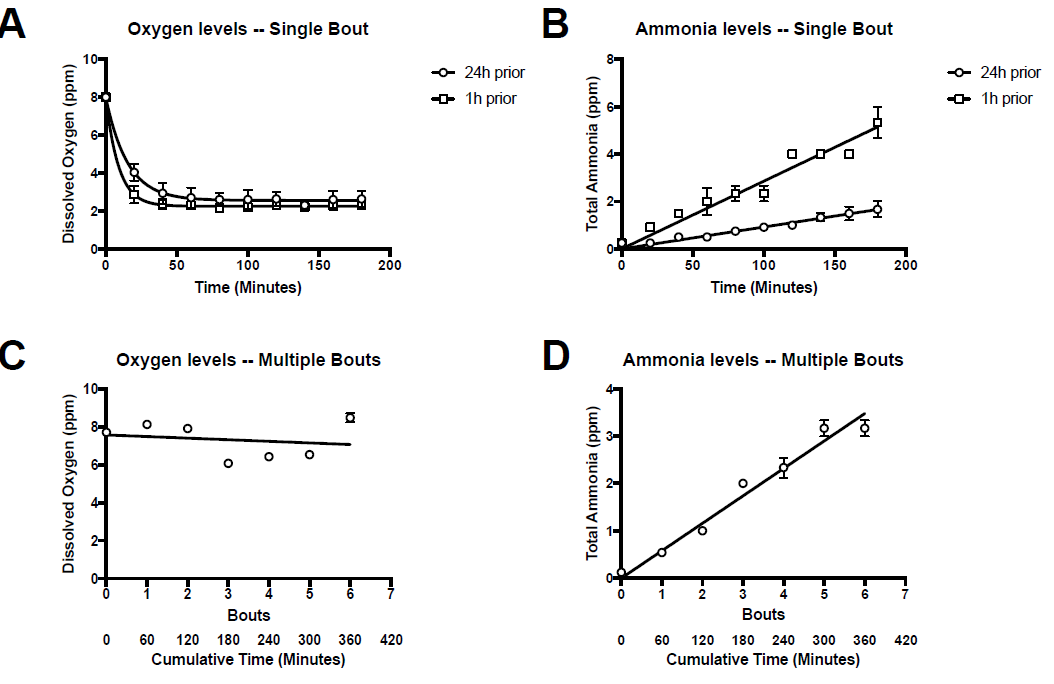
Water quality kinetics during immersion in the screening plate. (A) Dissolved oxygen over time within a single bout of immersion. Fish were fed 24h (black circles) or 1h (white squares) prior to experimentation. (B) Total ammonia over time within a single bout of immersion. Water was sampled from the same experiment as in (A). (C) Dissolved oxygen over multiple bouts of immersion. Fish were immersed for 60min/bout, and levels were measured prior to each bout. (D) Total ammonia over multiple bouts of immersion. Water was sampled from the same experiment as in (C). n=3 for all experiments; please see Tables 1 and 2 for curve fits.

Chemical libraries are typically constructed with compounds possessing a high degree of drug stability, putting forth the potential to re-use pre-filled screening plates if water quality is maintained over multiple dosing bouts. Thus, we next examined changes in water quality over multiple days of immersion, reusing the same water for each bout of immersion (60min/bout for 7 days; fish were fed 24h prior to the first bout, and not fed after). We found that ammonia concentration increased linearly with the number of dosing bouts/cumulative amount of dosing time (Fig 3C). The effective rate of total ammonia accumulation over multiple dosing bouts was 0.58±0.02ppm/hr, similar to the rate of 0.55±0.02ppm/hr observed in fish within a single dosing bout (i.e., in Fig 3A). This suggests that the introduction of fish into the wells itself does not stimulate appreciable ammonia excretion. Oxygen levels prior to each bout of immersion were similar each day (Fig 3D), with linear regression analysis revealing that the slope was not significantly different from zero (p=0.68). This suggests that the water became reoxygenated to baseline levels during storage. Due to this reoxygenation over multiple dosing bouts, the number of times in which the same drug/water volume may be reused is dictated by accumulation in uniodized ammonia (assuming the drug remains stable during reuse).

### System Validation

To test the fidelity of our system in detecting compounds affecting an adult physiology, we employed tail fin regeneration as a model system, and assessed the potential to recover known pharmacological inhibitors of regrowth in this process. Cyclopamine is a hedgehog pathway antagonist that modulates smoothened activity. Quint et al. previously demonstrated that chronic administration of 10µM cyclopamine starting at 2dpa results in impaired regrowth starting by 4dpa, and which is sustained to 7dpa ^19^. Lee et al. demonstrated that administration of 50µM cyclopamine at 4dpa was sufficient to reduce blastemal proliferation by 5dpa ^20^.

In our studies, we surmised that an immersion time of 60min would be tolerated in fish withheld from feeding 24h prior to dosing based on the fact that dissolved oxygen levels were sustained >3mg/L for the majority of this period. We employed a dosing regimen consisting of one hour of dosing in the 6-well plates per day starting on 1 day post amputation (dpa) and ending on 7dpa. Drug/water volumes were made fresh on each day of dosing. Re-epithelialization of the amputation stump via the formation of several layers of epithelial cells is observed by 1dpa ^21^; by waiting to administer compounds until this time point, we minimized the potential for compound administration to occur via wound entry. We compared fin regrowth in a) undosed free-swimming fish housed in standard 1.8L polycarbonate tanks, b) fish housed in the inserts and administered vehicles (DMSO or ethanol) via the 6-well screening plates, and c) fish housed in the inserts and administered different concentrations of cyclopamine (10µM, 50µM, or 100µM) via the 6-well screening plates (Fig 4). By 8dpa, we found that free-swimming fish housed in standard 1.8L polycarbonate tanks had similar fin regrowth compared to DMSO- and ethanol-treated fish housed in the screening system. Fish administered cyclopamine exhibited a dose-dependent reduction in fin regrowth, with significant differences observed for 50µM and 100µM cyclopamine, but not 10µM (Fig 4A).

**Figure 4:**
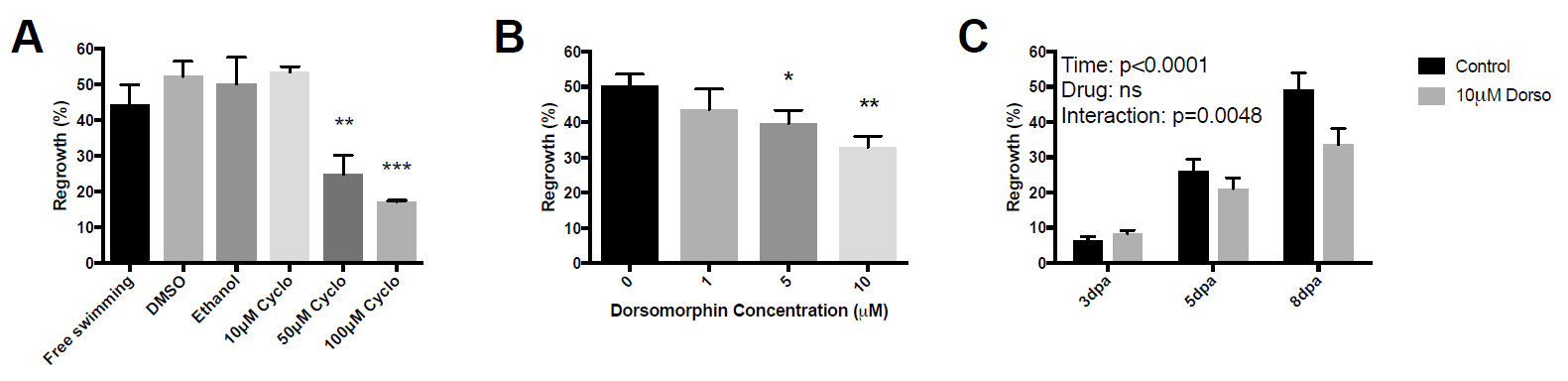
Validation of system fidelity in detecting chemical inhibitors of fin regeneration. For all studies, fish were administered drugs daily for 60min/day starting 1 day post amputation (dpa). A) % Fin regrowth at 8dpa in fish administered Cycloplamine. n=3 for all groups. (B) % Fin regrowth at 8dpa in fish administered Dorsomorphin. n=6 for all groups (C) Time-dependent effects of inhibition of fin regrowth in fish administered 10μM Dorsomorphin. n=5-6/group. For all studies, ^*^= p<0.05, ^**^= p<0.01, and ^***^= p<0.001.

Next, we assessed the potential to identify a predicted small molecule inhibitor of fin regeneration representative of lead compounds encompassed in small molecule libraries. DMH1 and LDN-193189 are optimized derivatives of dorsomorphin, a synthetic small molecule inhibitor of type I BMP receptors that was previously identified in an *in vivo* chemical screen in embryonic zebrafish ^22^. Thorimbert et al. previously showed that chronically administering 10µM of the BMP inhibitor DMH1 to adult zebrafish inhibits late-but not early-stage regrowth following amputation ^23^. Stewart et al. demonstrated similar inhibitory effects on late-but not early-stage regrowth following chronic administration of 5µM of the BMP inhibitor LDN-193189 ^24^. Due to off target effects, the concentrations in which dorsomorphin is active (4µM) and lethal (20µM) in zebrafish embryos are close in magnitude, providing a narrow window upon which to identify effective concentrations. Using the same dosing regimen as that used for cyclopamine studies, we compared fin regrowth in fish administered 1µM, 5µM, or 10µM dorsomorphin. By 8dpa, fish administered dorsomorphin exhibited a dose-dependent reduction in fin regrowth, with significant differences observed for 5µM and 10µM cyclopamine, but not 1µM (Fig 4B). Previous studies examining the effects of dorsomorphin derivatives (DMH1 and LDN-193189) on fin regeneration indicated that drug effects were manifested at later stages of regrowth. When we assessed whether effects on regrowth observed at 8dpa were observed at earlier time points, we found no evidence of regrowth deficits at 3dpa and 5dpa in fish treated with 10µM dorsomorphin (Fig 4C), with a two-way ANOVA revealing a significant effects of time (p<0.0001), drug (p<0.05), and time:drug interaction (p<0.05).

## DISCUSSION

The application of zebrafish chemical screens in post-embryonic physiologies has been limited by the lack of methods enabling rapid and cost-effective compound administration in adult animals. We have demonstrated the potential to integrate intermittent drug dosing with a dual-compartment chemical screening system to decrease the quantity of compounds required for screening by tenfold compared to the previous chemical screens in adult zebrafish ^11^. Inserts, lids, holders, and screening plates can be fabricated inexpensively using commercial desktop 3D printers. Further, the system design and dimensions can be readily modified to suit zebrafish of different developmental stages, as well as other aquatic species of different body shapes and sizes. We have included all CAD and STL files as supplementary information.

We characterized the kinetics of water quality to enable rational design of regimens for intermittent low-volume drug dosing. Our studies suggest that water quality kinetics dictate the design of dosing regimens in two ways. Over a single bout of dosing, the maximum time in which a fish may be immersed in the screening plate is dictated by the decrease in dissolved oxygen. The decay in dissolved oxygen can be altered through feeding, and is fully reversible after removal of the fish from the screening plate. In our studies, we surmised that an immersion time of 60min would be tolerated in fish withheld from feeding 24h prior to dosing based on the fact that dissolved oxygen levels were sustained >3mg/L for the majority of this period. Consistent with this notion, over the course of all experiments we observed 100% survival in control fish and near 100% survival in drug-administered fish, with only a single mortality occurring in a fish administered the highest dose of dorsomorphin, likely due to non-specific drug effects. Over multiple bouts of administration, the maximum allowable times in which a drug/water volume may be reused is dictated by the accumulation in uniodized ammonia. For the water temperature and pH employed in this study, sublethal poisoning from uniodized ammonia can occur at total ammonia levels of 2ppm. In our studies, the rate of increase in total ammonia was ~0.6ppm/hour; this rate was invariant whether the fish was subjected to a single long dosing bout, or multiple shorter bouts. This suggests each volume of water/drug may be used for a total of 2/0.6=3.3hr of cumulative dosing time. While we used fresh drug/water for each day of treatment, our studies suggest that repeated dosing of fish within the same drug/water volume could be a viable option to further conserve drug quantities during chemical screening.

To test the fidelity of the system, we assessed the potential to identify pharmacological inhibitors of adult tail fin regeneration. We found that intermittent dosing of cyclopamine mirrored effects of chronic cyclopamine exposure reported by Quint et al. in inhibiting later stages of regrowth in a dose-dependent manner, though higher concentrations were required to achieve an active concentration. In contrast, intermittent dosing of dorsomorphin replicated dose- and time-dependent effects of two dorsomorphin analogs, LDN-193189 and DMH1, at concentrations close to those previously reported to be active during tail fin regeneration. Following fin amputation, ectopic sonic hedgehog expression leads to bone fusion through a process mediated by activation of Bmp signaling ^21^, suggesting that Bmp signaling in the fin may be under the influence of the Shh pathway. This relationship does not appear to be reciprocal, as inhibition of bmp does not influence Shh signaling ^25^. The fact that dorsomorphin phenocopied the time-dependent effects of cyclopamine in inhibiting late regrowth reinforces a potential connection between Shh and Bmp type I receptor signaling in mediating fin outgrowth, and warrants further investigation in future studies.

Some limitations of our studies should be considered. First, we only assessed a single duration of dosing (1hr/day), and it is unclear to what degree altering this duration may affect assay sensitivity. While systematic investigations of the kinetics of drug absorption in zebrafish have yet to be performed, in goldfish, it has been proposed that the dosing time necessary for the fish to absorb a quantify of drug from a drug bath of concentration [C] scales as ~1/[C] ^26^. In this case, dosing fish for twice as long would require half the concentration to achieve the same quantity of drug to be absorbed; conversely, dosing fish for half as long would require twice the concentration. This suggests that different dosing strategies may be employed as different means to achieve the same amount of drug absorption: lower drug concentrations with a longer dosing time may be employed to conserve drug quantities; higher drug concentrations with shorter dosing times may be used to increase the throughput of drug administration, though at a higher monetary cost. While dissolved oxygen decay limits the duration in which fish may be immersed within a single loading bout, it is possible that dosing fish within a high-oxygen environment may reduce or even eliminate dissolved oxygen decay, and thus, extend the range of allowable dosing times. Third, while our studies of water quality kinetics suggest the potential to repeatedly use drug/water volumes over multiple dosing bouts, several practical issues need to be resolved before such strategies can be employed. This includes characterizing drug stability over multiple days, as well as potential loss of drug over multiple days as it is absorbed by the animals. Finally, it is noteworthy that drug actions can differ substantially depending on whether they are administered chronically or intermittently. While chronic administration is precluded in our system, intermittent administration more closely resembles drug kinetics and bioavailability associated with intermittent oral dosing in mammals ^27^, and thus has potential to enhance translational value in therapeutic screening applications.

In conclusion, we have developed a system for rapid and cost-effective chemical administration in adult zebrafish, expanding the potential for small molecule discovery in post-embryonic models of development, disease, and regeneration.

## ACKNOWLEDGEMENTS

Research reported in this publication was supported by the National Institute of Arthritis and Musculoskeletal and Skin Diseases (NIAMS) of the National Institutes of Health (NIH) under Award Number AR066061. The content is solely the responsibility of the authors and does not necessarily represent the official views of the National Institutes of Health. RYK would also like to acknowledge support from UW Royalty Research Fund Grant A88052, the University of Washington Department of Orthopedics and Sports Medicine.

